# Injectable Nanoporous Microgels Generate Vascularized Constructs and Support Bone Regeneration in Critical-sized Defects

**DOI:** 10.1101/2021.11.05.467366

**Authors:** Matthew D. Patrick, Jeremy F. Keys, Harshini Sureshkumar, Ramkumar T. Annamalai

## Abstract

Large and aberrant bone fractures require ossification and concomitant vascularization for proper healing. Evidence indicates that osteogenesis and vessel growth are coupled in bone fractures. Although the synergistic role of endothelial cells has been recognized, vascularizing large bone grafts remains a challenge and has apprehended the clinical translation of engineered bone constructs. Here, we describe a facile method to fabricate vascularized constructs using chitosan and gelatin-based microgels that promote osteogenesis of human mesenchymal stromal cells (MSC) while supporting endothelial sprouting and network formation. The microgels are enzymatically degradable and had a high hydration rate with a volume swelling ratio of ~560% and a polymer density of ~430 mg/cm^3^, which is comparable to that of native skeletal tissues. AFM indentation of the surface showed an average Young’s modulus of 189 kPa, falling in a range that is conducive to both osteogenesis and vasculogenesis. The osteogenic microgel containing chitosan, gelatin, and hydroxyapatite, mimicking the bone matrix, supported robust attachment, proliferation, and differentiation of MSC. On the other hand, the vasculogenic microgels containing only gelatin, enriched endothelial phenotype and enabled vascular networks formation when embedded in 3D matrices. Combining the two types of microgels created a hybrid construct that sustained the functions of both osteogenic and vasculogenic microgels and enhanced one another. Using a murine model, we also show that the osteogenic microgels regenerate bone in a critical-sized defect with >95% defect closure by week 12. These multifunctional microgels can be administered minimally invasively and can conformally fill large bone defects. This work lays the foundation to establish principles of designing multiphasic scaffolds with tissue-specific biophysical and biochemical properties for regenerating vascularized and interfacial tissues.

## 1. Introduction

Bone can heal without fibrotic scar formation by recapitulating the basic steps involved in fetal bone development during the healing process^1^. Despite its tremendous regenerative ability, large and aberrant bone fractures can result in slow, incomplete, or improper healing. An estimated 5-10% of all fractures will result in nonunion and develop pseudarthrosis^2^. There is an unmet clinical need to address these pathological conditions caused by traumatic injuries and critical bone loss due to tumor resection and failed arthroplasties. In clinical practice, a range of graft materials is used to treat nonunions, including autologous and allogeneic bone grafts, demineralized bone matrices, and a wide range of synthetic bone substitutes. Traditionally, autologous grafts have been the safest surgical reconstruction procedure and are efficacious and fast healing^3^. They boast the minimum risk of rejection but can result in intraoperative blood loss and donor-site morbidity. Moreover, creating a second surgical site results in a higher potential for infection and increases patient discomfort^4^. In some cases, an autograft isn’t possible due to the lack of a suitable donor site or the inability to conform the harvested graft to the defect shape^5^. In these situations, an allogenic graft is typically the next possible solution. An allograft eliminates the need for a secondary surgical site but has the potential for host rejection and disease transmission. Demineralized and decellularized allogeneic bone grafts, on the other hand, have a lower osteogenic potential due to their usual lack of cells and can result in high rates of bone resorption^6^.

In recent decades, tissue engineering approaches have shown promise in healing recalcitrant bone defects and raised considerable interest clinically and commercially. Despite the rapid advances in the field, commercially available products for orthopedic tissue regeneration are limited. The current tissue engineering methods and prostheses require extensive surgical procedures, resulting in long recovery times and further hindered by infection or loss of vascularization. For bone tissue engineering technologies to be feasible, it is essential to utilize the natural power of osteopotent cells, either by recruiting them from endogenous sources or delivering exogenous cells. Depending on the injury state, endogenous cell recruitment could be challenging, while exogenous cell delivery provides a more reliable and predictable outcome. Regardless of the source, the cells require an environment conducive to osteogenesis and neovascularization^7^. Consequently, the delivery of progenitor cells, along with an appropriate microenvironment, has the possibility of producing promising outcomes even under demanding circumstances.

Strategies that combine cells with osteoinductive materials have already seen clinical application^8^. However, they primarily use pre-formed scaffolds that require invasive surgery and accompany a plethora of complications^9, 10^. A minimally-invasive alternative would be suitable for reducing complications including infection, prolonged recovery time, and overall patient discomfort^11^. Studies have investigated the use of injectable polymer systems that form an insoluble hydrogel under physiological conditions^12^. These approaches are usually minimally invasive, but they lack the appropriate mechanics beneficial for ossification^13^. Although it can enhance the matrix mechanics, *in situ* chemical crosslinking of these polymers often induces cytotoxicity^11^. This constraint can be addressed by creating injectable cell-laden modules such as microcarriers or microcapsules out of the polymers with more appropriate mechanical properties. They can be fabricated, processed, and crosslinked to attain the desired mechanics before cells are seeded or implanted in the body to reduce cytotoxicity. Due to their microscale size range, they retain their injectability while providing a microenvironment for the cells^14^. Further, they can allow precise placement of different cell types and achieve organ-like cell densities that are not possible in a pre-formed scaffold^10^.

The use of adult stem cells, especially mesenchymal stem cells (MSC), has been widely studied in the context of bone healing due to their regeneration potential and wide availability from autologous tissue sources. Several *in vitro* and *in vivo* studies in small and large animal models have demonstrated the osteogenic potential of MSC. As of Oct 2021, more than 1340 clinical trials involving MSC have been registered in the ClinicalTrials.gov database for a wide range of diseases, at least 169 of them involving bone conditions. It is still argued whether the primary contribution of MSC is through morphogenesis and integration (direct effects) or through paracrine signaling (bystander effects)^15^. Also, there are some concerns about the long-term survival of the implanted MSC inside the host. But clinical studies show that delivering MSC in a material-based system enhances its bone regenerative potential^16^. Further, the physical characteristics of the materials contribute a great deal in determining the fate of the MSC^17, 18^. The mere geometry of the constructs can drive the lineage specification of MSC^19^. Together, scaffold design and fabrication can be a powerful tool in harnessing the therapeutic potential of MSC for bone defects.

Porous and nonporous polymeric microcarriers/microspheres such as Cultispher^®^ and Cytodex^®^ are commonly used as cell carriers. They generally have large solid fractions and low hydraulic permeability and hence do not emulate the high-water content of the native tissues^20^. To maintain a cytocompatible microenvironment and withstand the shear force during injections, hydration is a critical factor. At large solid fractions, the mechanics of polymeric scaffolds increasingly become more elastic and less viscoelastic^21^. Also, there is desirability for novel biomaterials and scaffold geometry rather than their feasibility and functionality^22^. Here, we describe a facile method to fabricate injectable microgels tailored to support osteogenic lineage commitment of MSC while supporting endothelial network formation. The osteogenic microgels tailored for MSC were fabricated with chitosan, gelatin, and hydroxyapatite to emulate the native bone structure. Chitosan is a natural polysaccharide resembling glycosaminoglycans of the bone^23^. Gelatin is a natural polypeptide^24^ that adds cell-binding domains and strength to the microgels^25^. Hydroxyapatite is the inorganic component of bone that adds to the rigidity of the matrix^26^. The vasculogenic microgels, on the other hand, tailored for endothelial cells (EC) and fibroblasts (FB) were fabricated with only gelatin. The amalgamation of these microgels can conformally fill irregularly shaped defects while enhancing and accelerating bone growth and vascularization.

In this study, we create off-the-shelf nanoporous injectable microgels and characterize them using clinically relevant primary human cell lines. We validate their injectability and other biophysical properties relevant for bone tissue engineering. We show that the osteogenic microgels support attachment, proliferation, and differentiation of bone marrow-derive MSC while the vasculogenic microgels enrich endothelial cells and promote vessel sprouting in a 3D matrix. Finally, we use a murine critical-sized defect model to show the effectiveness of the osteogenic microgel in regenerating calvarial bone. Together, the microgels possess physical and biochemical properties conducive for regenerating vascularized bone in critical-size defects.

## 2. Materials and methods

### 2.1. Cells and biopolymers

All studies involving human and animals cells or tissues are conducted according to approved protocols by the Institutional Biosafety Committee (IBC) protocol and Institutional Animal Care and Use Committee (IACUC) at the University of Kentucky, respectively. For fabricating osteogenic microgels, human bone marrow-derived mesenchymal stem cells (MSC) from 4 normal healthy adult donors (a 26-year-old female, a 20-year old female, a 23-year-old male, and a 19-year-old female) were obtained from a commercial vendor (RoosterBio) and screened for viability, proliferation, and differentiation potential. Since no significant difference was noticed, MSC from a single donor (a 26-year old female) with a population density level (PDL) of 9-20 was used for the rest of our studies. For the expansion of MSC, growth media made from RoosterBasal^™^-MSC and supplemented with RoosterBooster^™^-MSC was used. Dulbecco’s Modified Eagle’s Medium (DMEM, Gibco) with 10% FBS (Gibco) with or without the osteogenic supplements were used for all experiments. To induce osteogenic differentiation of MSC, DMEM was supplemented with 10% FBS, 10 mM β-glycerol phosphate disodium (β-G2P, Sigma), 0.2 mM L-ascorbic acid 2-phosphate (Sigma) and, 100 nM dexamethasone (Sigma).

The vasculogenic microgels were seeded with human lung fibroblasts (FB, Lonza) from a 29-year-old male donor and human umbilical vein endothelial cells (EC, Lonza) from three Caucasian male infants were obtained and screened for proliferation, viability, and sprouting. One of the EC donor lots was used for all studies. FB were expanded using DMEM with 10% FBS, while EC were expanded using VascuLife^®^ VEGF endothelial medium complete Kit (Lifeline Cell Technology). EC and FB with PDL <20 were used for all our studies. For coculture experiments, VascuLife^®^ VEGF endothelial medium was used. For mice implantation studies, murine adipose-derived mesenchymal stromal cells (MSC) were isolated from inguinal fat pads of adult C57BL/6J mice^14, 27^. These cells have the established potential to regenerate bone^28^. Briefly, the fat pads were isolated, washed in serial dilutions of betadine, minced, and the matrix was degraded in 0.1 wt% collagenase solution. The resulting single-cell suspension was cultured in DMEM with 10% FBS, and adherent cells were isolated, passaged (5^th^), and used for experiments. Cells were seeded on microgels by adding the 100 μL of cell suspension 0.25-0.5 million cells onto microgels suspended in growth media. Then the suspension was kept in the incubator for 1 hour to allow cell attachment to the surface of the microgels. During this period, the suspension was gently mixed every 10 min to prevent aggregation and allow uniform seeding. Finally, the microgels were washed with fresh media to remove unbound cells, suspended in 1 mL of fresh media, and kept in the incubator until further treatment.

The biopolymers used for fabricating microgels include gelatin from porcine skin (type A, 175 bloom, Sigma), low molecular weight chitosan (75-85% deacetylated, 50,000-190,000 Da, Sigma), and fibrinogen from bovine plasma (Type I-S, ≥75% clottable protein, Sigma). A 6-8 wt% gelatin stock solution was made by dissolving lyophilized gelatin in calcium and magnesium-free phosphate-buffered saline (10 mM PBS; Invitrogen). The suspension was kept at 40°C until the gelatin was completely dissolved. Similarly, a 1 wt% chitosan stock solution was made by dissolving the 0.25 g of chitosan flakes in 25 mL of 0.02 N acetic acid and stirring for seven days at 4°C. The solution was then centrifuged at 10,000 g to remove undissolved debris and blended with a suitable amount of gelatin stock for making microgels. For formulations containing hydroxyapatite (HA), a 1 wt% HA in 10 mM PBS stock (nano grade, <200 nm particle size, Sigma) was made and sonicated for 5 min before use.

### 2.2. Microgel fabrication

The microgels were fabricated using a simple water-in-oil emulsion technique, as shown^14, 29^. Briefly, a 1-5 mL of solubilized gelatin solution was dispersed dropwise into a 90 ml of stirred polydimethylsiloxane (PDMS, viscosity = 100 cS, Clearco Products Co., Inc.) bath kept at 40°C. The PDMS-gelatin mixture was mixed continuously with an impeller at desired rpm for 5 min. Then the temperature of the PDMS bath was reduced using ice to promote gelation of the gelatin, while the emulsion was continually stirred for an additional 30 min. The mixture is then transferred to 50 mL centrifugal tubes and centrifuged at 500 g for 5 min to separate the PDMS from gelatin. After centrifugation, the bulk PDMS supernatant was decanted without disturbing the pelleted microgels. The residual PDMS was thoroughly removed by subsequent washes with 10 mM PBS containing 1% TWEEN 20 (Sigma). Finally, the isolated gelatin microgel spheres were crosslinked to improve stability by suspending them in a 1% genipin (Wako Chemicals) in 10 mM PBS solution for up to 48 hours. Then they were washed in 100% ethanol to remove unbound genipin. The crosslinked microgels were then sorted based on the size rages using nylon filters and stored at 4-8°C until further use.

Osteogenic microgels were made by emulsifying a blended mixture with a final concentration of 6 wt% gelatin, 0.25-0.75 wt% chitosan, and 1 wt% HA. Vasculogenic microgels were made by emulsifying 6 wt% gelatin solutions. For vasculogenic assays, the microgels were embedded in bulk fibrin hydrogel 2.5 mg/mL in serum-free DMEM. For murine implants, 2.5 mg/mL fibrin was used as a carrier gel to concentrate and contain the microgel within the defect.

### 2.3. Mass and volume swelling ratios of microgels

The volume swelling ratio (Q) of the microgels was calculated by measuring the size and volume of hydrated and dehydrated microgels, as shown previously^30^. The value of Q was calculated using the formula:

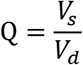

where *V_s_* is the volume of the microgel swollen under equilibrium hydration conditions, and *V_d_* is the volume of the dry microgels. After crosslinking, the microgels were thoroughly washed in 100% ethanol followed by PBS and suspended in fresh PBS overnight at room temperature. Bright-field images of the microgels were taken the next day to measure the diameter and volume of the hydrated microgels. Then they were flash-frozen in liquid nitrogen and lyophilized overnight. The size of the dried microgels was measure using bright-field imaging. Finally, the polymer density (*ρ_p_*) was calculated using the formula:

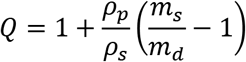

where *m_s_* and *m_d_* are the mass of swollen and dried microgels, respectively, and *ρ_s_* is the density of the solvent (10 mM PBS).

### 2.4. Cell proliferation and viability

The cell proliferation was analyzed by quantifying the total DNA content of samples using the Quant-iT^™^ PicoGreen^™^ dsDNA Assay Kit (ThermoFisher). Lamda DNA standard was used for generating the calibration curve. Briefly, samples were collected at specific time points and homogenized in 1X Tris-EDTA buffer (200mM Tris-HCl, 20 mM EDTA, pH 7.5), and centrifuged at 10,000 g for 10 min to remove cell debris. The supernatant was then mixed with PicoGreen buffer, incubated in the dark for 5 min, and the fluorescence was read at Ex/Em wavelength of 485/528 nm^31^.

The viability of cells was assessed using a LIVE/DEAD cell Imaging Kit (Invitrogen) following the manufacturer’s instructions. Briefly, adherent cells were washed with 10 mM PBS and stained with 2 μM calcein AM, and 4 μM ethidium homodimer-1 in 10 mM PBS and incubated for 30 min at room temperature. After incubation, the samples were imaged fluorometrically using a Nikon A1R confocal microscope. Live and dead cells were counted using Fiji (ImageJ) software (National Institutes of Health), and the ratio of live to dead was calculated.

### 2.5. Orthocresolphthalein complexone (OCPC) assay for calcium

The *o*-cresolphthalein complex one (OCPC) method was used to quantify calcium deposition by the MSC in microgels as previously described^14^. Briefly, the samples were homogenized in 1X Tris-EDTA buffer and centrifuged at 10,000 g for 10 min. The supernatant was collected and mixed with a working solution containing 5% ethanolamine/boric acid buffer (Sigma), 5% o-cresolphthalein (Sigma), and 2% hydroxyquinoline (Sigma) in distilled water. The mixture was incubated at room temperature in the dark for 10 min, and the absorbance was read at 575 nm. CaCl2 (Sigma) dissolved in water at different concentrations served as standards. Microgels with HA in the formulation were excluded from this assay.

### 2.6. Sircol^™^ assay for collagen

The collagen deposition by the MSC was quantified using a Sircol soluble collagen assay kit (Biocolor). Briefly, the samples were homogenized in 1X Tris-HCl buffer and centrifuged at 10,000 g for 10 min. The supernatant was collected, and a small volume was mixed with 1 mL of Sircol^™^ reagent, gently mixed for 30 min, and centrifuged at 10,000 g for 10 min. The supernatant was decanted, and a 750 μL of acid-salt wash reagent was added to the pellet, mixed, and centrifuged at 10,000 g for 10 min. The supernatant was decanted, and a 250 μL of alkali reagent was added to the pellet, vortexed, and incubated for 5 min at room temperature. The absorbance of the final solution was read at 555 nm. Rat-tail collagen standards were used for calibration.

### 2.7. Alkaline phosphatase (ALP) assay

ALP activity of samples containing MSC was quantified as an early-stage osteogenic marker. A commercially available ALP activity fluorometric assay kit (BioVision) was used according to manufacturer instructions. Briefly, the samples were homogenized in ALP buffer and centrifuged at 10,000 g for 10 min. A 100 μL of the supernatant was mixed with 20 μL of 0.5 mM 4-methylumbelliferyl phosphate (MUP) substrate and incubated at 25°C in the dark for 30 min. Then a 20 μL of stop solution was added to the mixture, and the fluorescence was measured at Ex/Em of 360/440 nm. Standards of 0.1-0.5 nmol of MUP substrate along with purified ALP enzyme were used for calibration.

### 2.8. RNA isolation and quantitative gene expression analysis

For gene expression analysis, the samples were collected in 500 μL of TRIzol^™^ reagent (Invitrogen) in a 1.5 mL vial, vortexed, and 100 μL of molecular grade chloroform (Sigma) was added to the tube and incubated at room temperature for 15 min. The samples were centrifuged at 10,000 g at 4°C for 10 min. The aqueous phase was transferred carefully without disturbing the interphase into a fresh 1.5 mL tube, and 10 μg of glycogen was added as a carrier. Then 500 μL of isopropanol (Sigma) was added to the aqueous phase and incubated for 10 more min at room temperature. The samples were centrifuged at 12,000 g at 4°C for 10 min. The supernatant was decanted, and to the pellet, 1 mL of 75% ethanol was added, vortexed, and centrifuged at 7500 g at 4°C for 5 mins. The resulting RNA pellet was air-dried, and 10-20 uL of RNAse free water was added and heated at 60°C for 10 min.

A SuperScript^™^ III Platinum^™^ One-Step qRT-PCR Kit (ThermoFisher) and TaqMan probes (ThermoFisher) for osteogenic genes were used to analyze gene expression profiles using QuantStudio 3 real-time PCR system (ThermoFisher). The TaqMan probes used for our studies are *ALP* (Hs01029144_m1), *BGLAP* (Hs01587814_g1), *COL1* (Hs00164004_m1), IBSP (Hs00173720_m1), *SP7* (Hs00541729_m1), *SPP1* (Hs00959010_m1), *RUNX2* (Hs01047973_m1), and *GAPDH* (Hs99999905_m1) was used as housekeeping control. Gene expression data were processed in MATLAB, and heatmaps and dendrograms were generated using the clustergram function of the bioinformatics toolbox.

### 2.9. Immunofluorescence staining and confocal microscopy

Samples were fixed in 10% aqueous buffered zinc formalin (Z-Fix, Anatech) for at least 6 hours for immunofluorescence staining. Phalloidin conjugates (AF488, ThermoFisher) were used to stain the F-actin of the cell cytoskeleton. For identifying MSC in co-cultures studies, the MSC were stained with CellTracker^™^ Green CMFDA (Invitrogen), and fibroblasts were stained with CellTracker^™^ Violet BMQC (Invitrogen) before seeding them in microgels. The endothelial sprouts were stained with an endothelial cell-specific marker Ulex Europaeus Agglutinin I (UEA-I, Vector Laboratories). DAPI (Invitrogen) was used as a nuclear counter-stain. Before staining, the fixed samples were washed and permeabilized with 0.1% Triton X-100 (Sigma) in 10 mM PBS. The samples were then stained for 1 hour in the dark, and z-stacks were acquired using a Nikon A1R confocal microscope. All the acquired optical sections were processed using Fiji (ImageJ) software.

### 2.10. Electron microscopy

For electron microscope imaging, fixed samples were dehydrated in 100% ethanol, flash-frozen, and lyophilized. The dried samples were then sputter-coated with a 5 nm thick layer of platinum using Leica ACE 600 sputter coater. The samples were then imaged using the FEI Quanta 250 environmental scanning electron microscope (SEM). To image the inner core of the microgel, a Helios SEM system equipped with a focused ion beam (FIB) was used. First, a section of the sample was coated with a 300 nm carbon layer to protect the specimen for the setup and then ablated to half the size (~40 μm deep) using the FIB. The samples were then imaged using the SEM.

### 2.11. AFM measurement and compression testing

A JPK-Bruker Nanowozard^™^ 4a atomic force microscope (AFM) was used to image the microgels as described^32^. qp-BioAC cantilever probe (Nanosensors) with rounded tip apex with a radius of curvature 30 nm, force constant 0.1 N/m, tip height 7 μm were used for Quantitative Imaging (QI) in the AFM. Data analysis was performed by using JPK Data Processing software. Force-displacement data was extracted, and Young’s moduli map of the specimen was constructed using a Hertzian model fit on the ‘extend’ curve with a parabolic profile for the tip shape^3333^. Briefly, the elastic modulus was calculated using the formula:

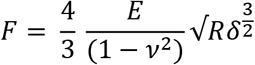

Where F is force, δ is displacement, R is the radius of curvature of the tip, E is Young’s modulus, and ν is the Poisson’s ratio of the specimen (assumed to be 0.5).

For compression testing, bulk hydrogels constructs were prepared in a 12-well flat-bottom plate with the same formulation as the microgels. They were then cross-linked using 1% genipin, washed in ethanol, and rehydrated in 10 mM PBS. Using biopsy punches, 6 mm disks were isolated and compression tested using Universal Testing Machine (Instron) at a strain rate of 10 mm/min. The elastic modulus was calculated from the linear region (between 5-15% strain region) of stress-strain curve.

### 2.12. Injectability assessment

To demonstrate the minimally invasive application of the microgels, they were suspended in PBS (~2 mg of microgel/mL), collected in a 1 mL syringe (BD Plastics), and injected through a 16-25 gauge hypodermic needle at a flow rate of 0.1 mL/s. The injection force was simultaneously measured using a Universal Testing Machine (Instron) equipped with a 10 N load cell. PBS without microgels served as control. Then to characterize the viability of the cell-laden microgels subjected to injection force, they were concentrated and injected through a 16 gauge needle and collected in a vial and characterized for cell viability. Viability was assessed at 1 and 24 hours using a live/dead cell imaging kit and counting using Fiji (ImageJ) software (National Institutes of Health).

### 2.13. Murine osteotomy model

All animal procedures were performed in compliance with the protocol approved by IACUC at the University of Kentucky. As per the ARRIVE guidelines^34^, we report the following essential details of the study. To validate the bone regeneration capacity of osteogenic microgels seeded with murine ASC, a well-established critical-sized calvarial defect model was used, as described^14^. For our pilot studies, 3-4-month-old adult C57BL/6 mice (nine male and one female) were used and randomly assigned to treatment groups. Briefly, mice were anesthetized (1.5 mg of ketamine/kg + 1.5 mg of xylazine/kg body mass of mice), a bland ophthalmic ointment was applied, and the surgical site was shaved and wiped with 70% ethanol, followed by 10% povidone-iodine solution. A 1.5-2 cm incision was made along the sagittal midline to expose the parietal lobe parallel to the lambda suture. Next, the periosteum was removed with a cotton swab, and a PiezoSurgery (Mectron) apparatus equipped with a 3 mm osteotomy insert (OT11) was used to create a 3 mm critical-sized defect in the parietal bone. After removing the 3 mm cortical bone, an appropriate construct was placed in the defect, and skin was sutured with 6-0 vicryl suture (Ethicon). Then, 1 mL of saline was administered intraperitoneally for hydration followed by subcutaneous administration of buprenorphine SR (1.5 mg/kg body mass) for analgesia and placed into a recovery cage and monitored as per the established post-operative animal care protocols. Animals were monitored daily for 7 days and weekly thereafter to ensure post-operative recovery.

### 2.14. *In vivo* microCT imaging and morphometric analysis

*In vivo* bone formation was assessed using longitudinal microcomputed tomography (microCT), performed using a high-resolution small animal imaging system SkyScan1276 microCT (Bruker, Billerica, MA) with an Al 1mm filter with the voltage set to 70 kV and image pixel size 40.77 μm. Images were then reconstructed (NRecon) with smoothing set to 3, ring artifact correction set to 4, and beam hardening correction set to 40%. For morphometric analysis, cylindrical phantoms (2 mm dia) with a precise bone mineral density of 0.25 and 0.75 g/cm^3^ placed in deionized water were scanned and reconstructed with identical settings. Reconstructed files were then imported to CTan software for morphometric and mineral density analysis. The contralateral bone was used to determine the greyscale threshold to use for cortical bone. A 1.25 x 0.32 mm^3^ (radius x height) cylindrical VOI was placed in the contralateral parietal bone, and the greyscale threshold was determined using a 5 level OTSU algorithm inside the VOI, level 3 was taken for as the cortical bone threshold. The defect was then analyzed using the same VOI and level 3 threshold. All ROI for analyses are restricted only to the defect site and any mineralized tissue formation in apposition to the native bone was not included in our analysis to enhance rigor.

### 2.15. Statistics

All measurements were performed at least in triplicate. Data are plotted as means with error bars representing the standard deviation. The Pearson correlation coefficient (r) was used to evaluate the linear correlation between two variables. Statistical comparisons were made using Student’s t-test (two-tailed and unequal variance), Mann-Whitney rank-sum test, Holm-Sidak test, and two-way ANOVA with a 95% confidence limit. Differences with p < 0.05 were considered statistically significant.

## 3. Results

### 3.1. Fabrication and characterization of nanoporous microgels

The emulsification of gelatin hydrogel solution in PDMS yielded spherical hydrogel droplets, which upon crosslinking with genipin, yielded insoluble nanoporous microgels. A microgel diameter range of 200 ± 50 μm under equilibrium hydration conditions was deemed suitable for our application. This specific microgel size range provides sufficient surface for cell attachment and proliferation while allowing minimally invasive delivery using a 16-18 gauge needle (inner diameter 800 −1200 μm). A parallel-blade and a propeller-type impeller were tested at different speeds to maximize the yield of microgels in the desired size range. A 6 wt% gelatin stock solution was used for the test, emulsified at 40°C for 5 min, and rapidly cooled using an ice bath. Post crosslinking with genipin for 24 hours, the microgels showed a size distribution that varied with speed and impeller type. With both the impellers, increasing the emulsification speed reduced the size distribution of microgels (**Fig.1A**). The parallel-blade impeller yielded microgels with sizes showing a bimodal distribution at 250 and 750 rpm. On the other hand, the propeller impeller yielded microgels with sizes showing normal distribution and a linear decrease in mean and median values with an increase in mixing speed (**Table 1, Fig.1A**). Further emulsification at 500 rpm generated the maximum number of microgels in our desired size range for a given load. Based on these results, a propeller impeller and 500 rpm mixing speed were deemed appropriate to create microgels for the rest of the study.

**Figure 1.**
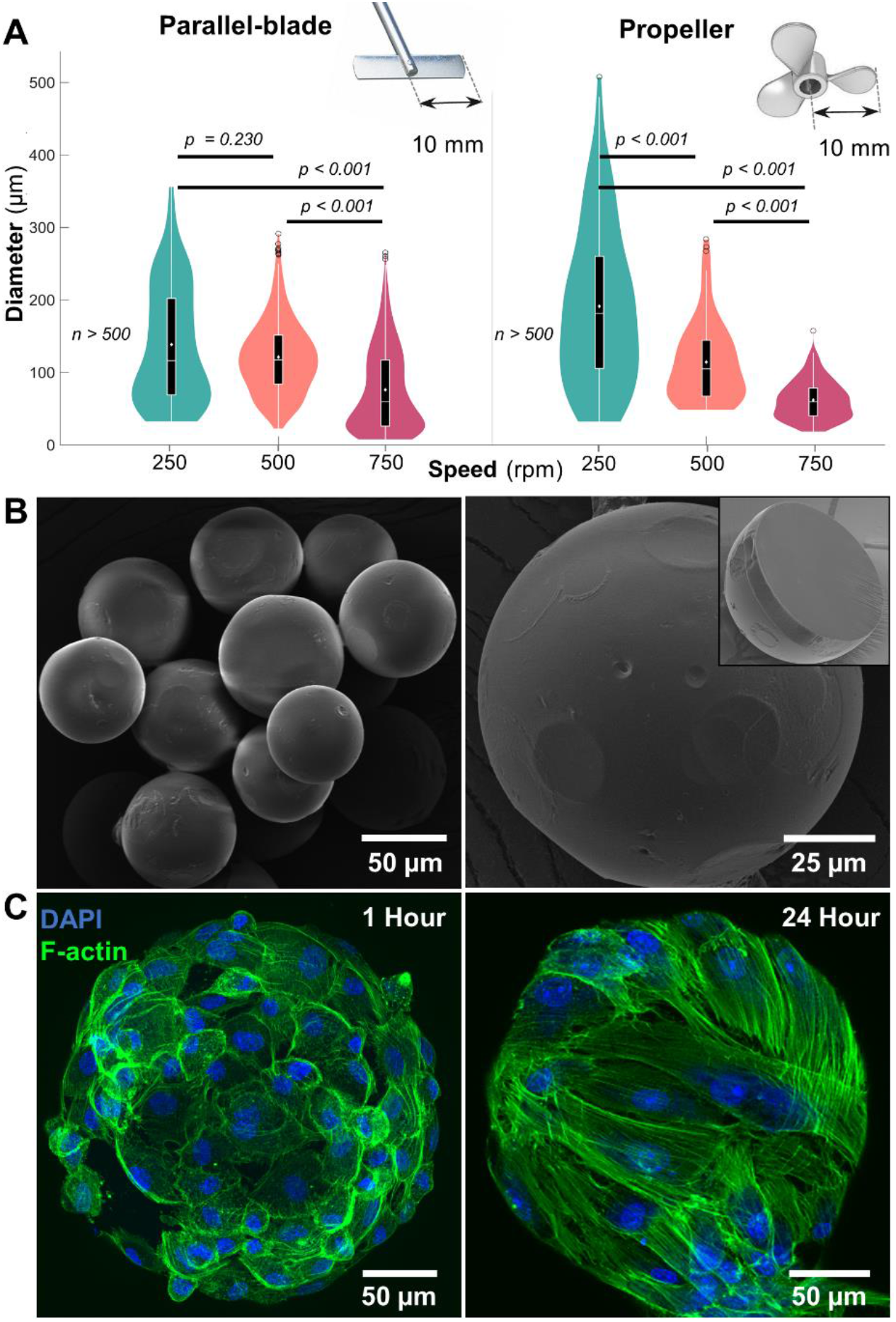
Microgel fabrication and characterization. **A)** Size distribution of gelatin microgels made using straight-blade and propeller impellers at mixing speeds of 250, 500, and 750 rpm (n ≥ 500). Violin plot: White dots and lines represent mean and median, respectively. Black boxes, whiskers, and circles represent interquartile ranges, 25/75 quartile ranges, and outliers, respectively. **B**) Scanning electron micrographs of dehydrated microgels with reduced size and a smooth outer surface. (Inset) Cross-section of a dehydrated microgel showing the core with a uniform, densely packed matrix. **C**) Confocal micrographs of hydrated gelatin microgels seeded with mesenchymal stem cells showing a stretched cytoskeleton within 24 hours.

**Table 1:** Size distribution of microgels

| | Speed (rpm) | Mean ($\mu\text{m}$) | Median ($\mu\text{m}$) |
| --- | --- | --- | --- |
| | 250 | 138.5±81 | 117.4 |
| Parallel plate | 500 | 121.2±51 | 117.4 |
|  | 750 | 76.1±58 | 59.6 |
| Propeller | 250 | 191.3±109 | 181.7 |
|  | 500 | 114.4±55 | 104.9 |
|  | 750 | 61.9±26 | 59.5 |

Scanning electron micrographs of dehydrated microgels showed a smooth outer surface (**Fig.1B**). Upon milling the surface using a focused ion beam (FIB) with nanometer precision, the inner matrix of a dehydrated microgel revelated a uniform densely packed structure (**Fig.1B** inset). To investigate the cytocompatibility of the microgels, human bone marrow-derived MSC were seeded on the surface, and samples were collected at 1 hour and 24 hours. When stained for the cytoskeletal molecule F-actin, the samples revealed that the cells were firmly attached to the surface within an hour of incubation (**Fig.1C** left). The cells continued to spread on the surface and engulfed the microgel within 24 hours (**Fig.1C** right).

The swelling behavior and the polymer density of the microgels were then analyzed by measuring the mass and volume ratios of the microgels when equilibrated in PBS at room temperature and after lyophilization. The lyophilized microgels rapidly absorbed water and exhibited a high degree of hydration (**Fig.2A**) with a volume swelling ratio of 567% and a polymer density of 431 ± 85 mg/cm^3^ in PBS (**Fig.2B**). The addition of other materials to the gelatin matrix, including 0.25% chitosan or 1% hydroxyapatite (HA), did not significantly affect their swelling behavior. However, the compressive modulus of the bulk hydrogels changed slightly depending on the material (**Fig.2C**). Quantitative imaging of a hydrated microgel using an atomic force microscope (AFM) showed a nanoporous surface with an average Young’s modulus of 189.9 kPa (**Fig.2D**).

**Figure 2.**
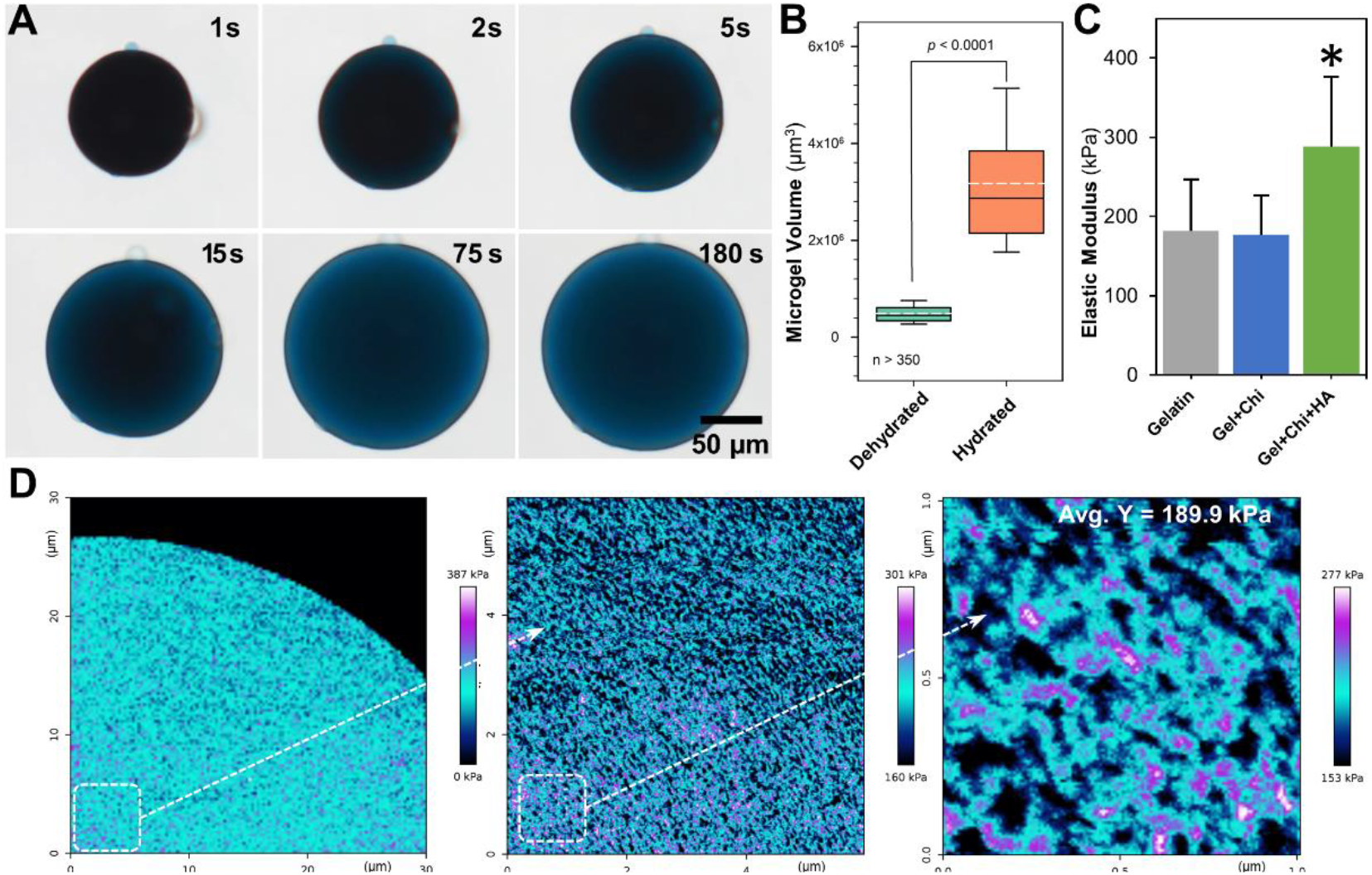
Microgel swelling and elasticity. **A)** Temporal bright-field images of a microgel being hydrated in PBS. **B)** The change in volume of the microgels at the equilibrium hydration condition (n ≈ 350). **C)** The compressive modulus of the gelatin hydrogels changed slightly depending on the formulation (n = 10). **D)** Young’s modulus map of a hydrated gelatin microgel at different magnifications. Micrographs were obtained using an AFM in QI mode. Young’s moduli map was constructed using a Hertzian model fit on the ‘extend’ curve with a parabolic profile for the tip shape.

### 3.2. Customization of microgel composition to mimic bone matrix

To enhance the osteogenicity of the microenvironment, 0.25-0.75 wt% chitosan (low MW) and 1 wt% of hydroxyapatite (HA, <200 nm size) were added to the microgel formulation. The HA concentration was limited to 1 wt% to mimic bone consistency without compromising the integrity of the microgels. Blending chitosan up to 0.75% with gelatin did not interfere with the thermogelation process and produced microgels with no significant change in size distribution (**Fig.3A, B**). Likewise, the addition of HA to the hydrogel at 1 wt% did not affect the fabrication process or microgel size distribution. But the HA aggregated to some extent within the microgel during fabrication. Compared to the smooth surface of the gelatin microgels (**Fig.3C**(i)), the addition of chitosan (**Fig.3C**(ii)) and chitosan+HA (**Fig.3C**(iii)) resulted in slight variations in the matrix texture. Regardless of the formulation, the microgels showed a uniform and densely packed core, indicating an even blending of materials. Although chitosan is highly biocompatible, they lack cell attachment domains. Therefore, for the subsequent studies, 0.25% chitosan was deemed suitable to promote maximal cell attachment on the surface.

**Figure 3.**
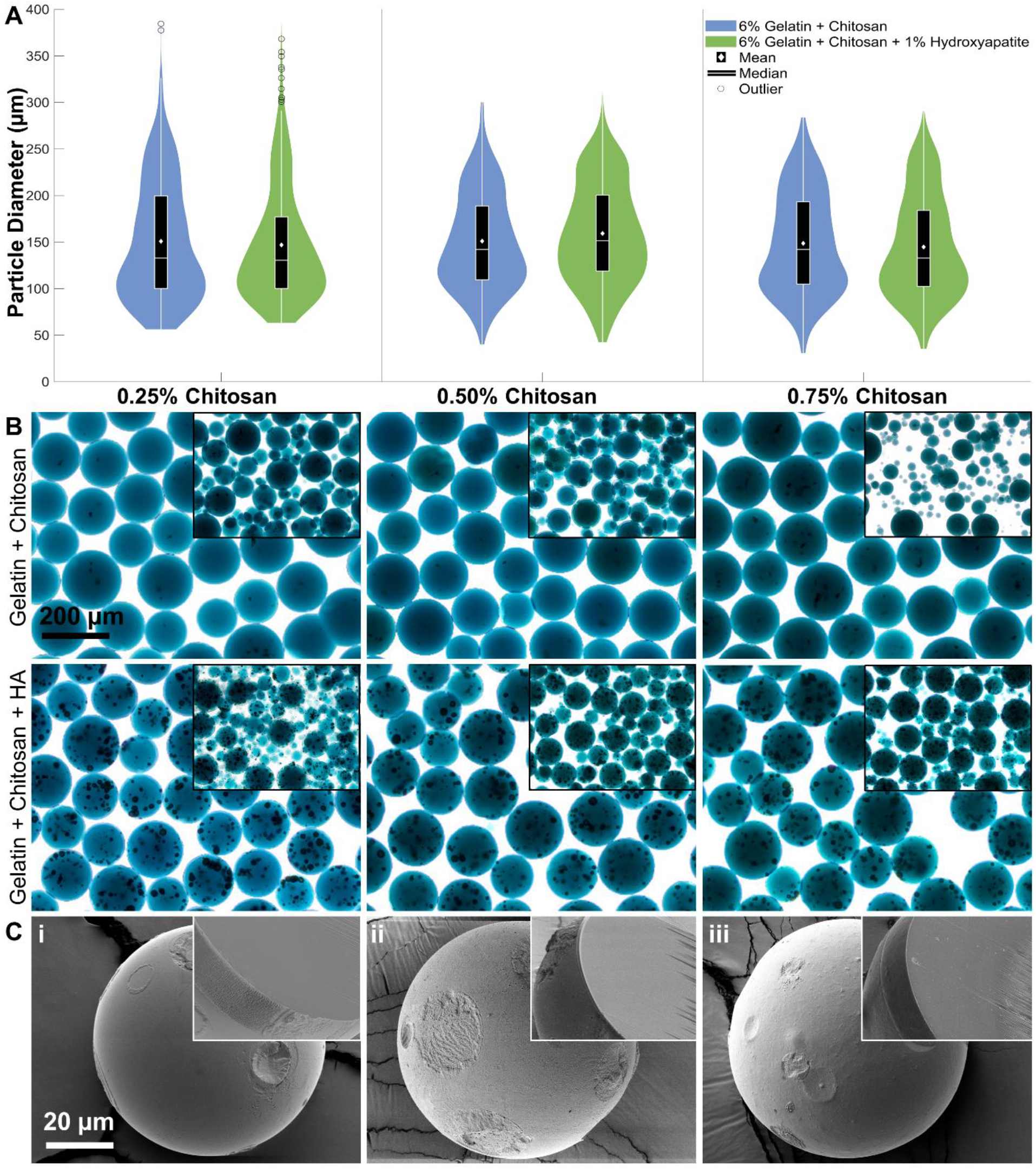
Osteogenic microgel fabrication and characterization. **A)** Size distribution of gelatin microgels fabrication after adding 0.25, 0.50, or 0.75 wt% chitosan and 1 wt% hydroxyapatite (n ≥ 100). **B)** Representative bright-field micrographs of the 150-250 μm microgels made from different chitosan and HA concentrations. The HA aggregates can be seen as dark spheroids within the microgels. The insets show the unsorted microgels of each formulation. ANOVA shows no statistical difference between groups. **C)** Scanning electron micrographs of dehydrated microgels formulations: i) Gelatin; ii) Gelatin+0.75 wt% chitosan; iii) Gelatin+0.75 wt% chitosan+1 wt% hydroxyapatite. (Inset) Cross-section of dehydrated microgels showing a uniform and densely packed matrix with slight variations in the texture between formulations.

### 3.3. MSC seeded on osteogenic microgels show enhanced osteogenesis

To validate the enhanced osteogenic potential of the osteogenic microgels, human bone marrow-derived MSC were seeded on the microgels. Samples from short-term and long-term cultures were collected and analyzed for cell attachment, surface coverage, and differentiation. Samples stained for cytoskeletal F-actin showed that MSC were firmly attached to the surface within an hour of seeding (**Fig. 4A**). Within 12 hours of seeding, the cells start to spread and cover the surface, and by 24 hours, the microgel surface is fully covered in cells (**Fig. 4A**). A one-way ANOVA statistical analysis shows no significance in the number of cells attached per sphere (*p* = *0.259*) or the surface fraction of microgel covered in cells over time (**Fig. 4B** top and middle bar graphs). However, there was a significant increase in cell spreading area by 12 and 24 hours compared to that of 1-hour in microgels with HA (**Fig. 4B** bottom bar graph). Overall, the addition of chitosan or HA did not alter MSC attachment or spreading and showed results similar to control gelatin microgels shown above.

**Figure 4.**
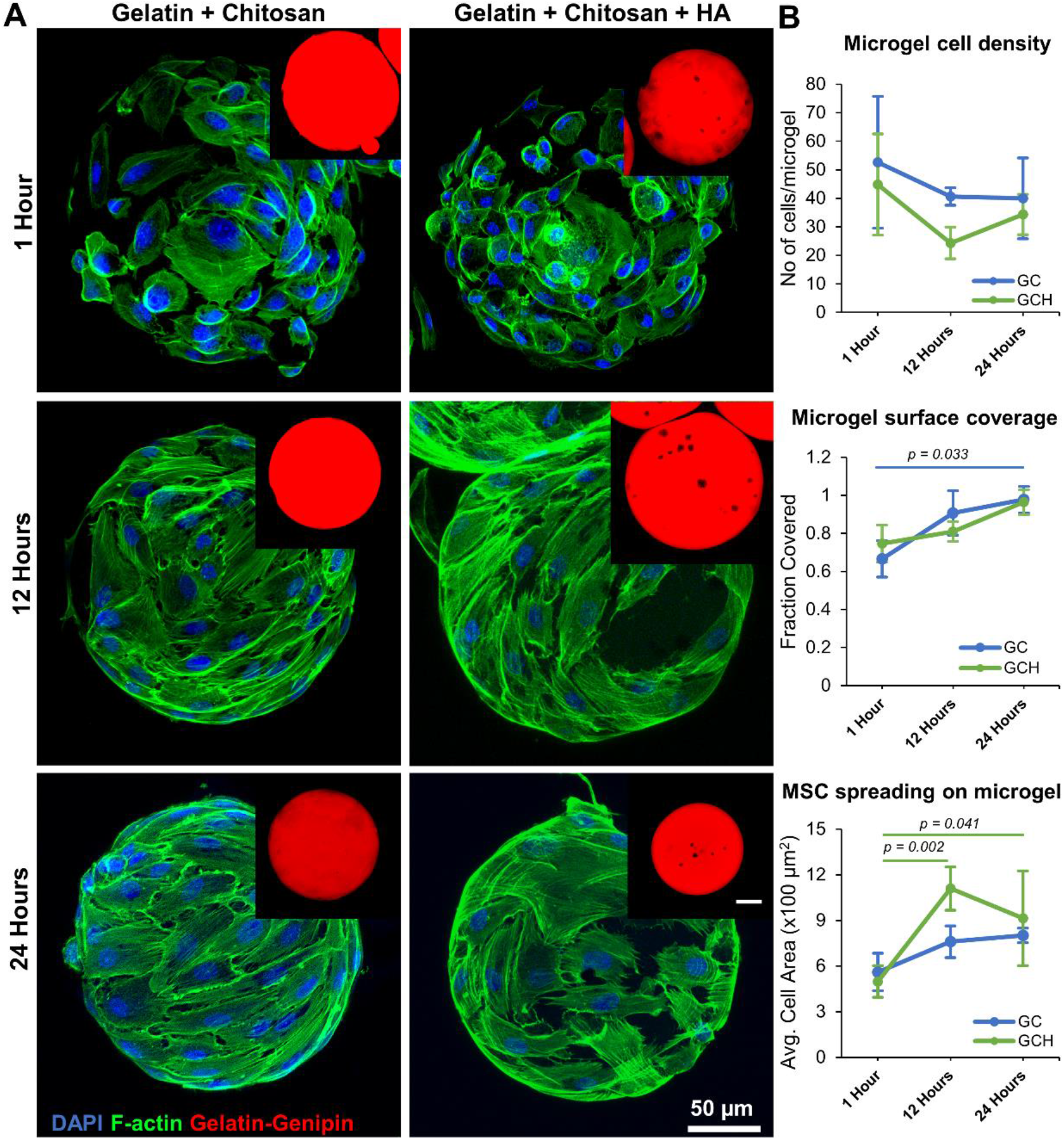
MSC seeded on osteogenic microgels. **A)** Confocal stacks of microgels seeded with MSC at different timepoints. The cells are stained for cytoskeletal F-actin using phalloidin (green) and nuclei with DAPI (blue). The insets show the red fluorescence of the gelatin matrix crosslinked with genipin. **B)** (top to bottom) the average number of MSC attached per microgel, the fraction of microgel surface covered by attached MSC, and the average area of the MSC attached to the microgel surface over time (n = 5). Bar chart legend: GC-Gelatin+0.25% Chitosan, GCH-Gelatin +0.25% Chitosan+1%HA.

To investigate the osteogenicity of the microgels, MSC-laden microgels were cultured for 3 weeks in growth and osteogenic conditions. The transcriptomic analysis of samples collected at 0, 7, 14, and 21 days showed that essential osteogenic genes including osteocalcin (*BGLAP*), bone sialoprotein (*IBSP*), collagen type I (*COL1A1*), Runt-related transcription factor 2 (*RUNX2*), and alkaline phosphatase (*ALP*) are upregulated in MSC (**Fig.5A**). Interestingly gene expression of MSC in growth conditions mirrored the osteogenic gene upregulation seen in osteogenic conditions. Likewise, changes in material composition influence osteogenic gene expression. MSC Proliferation, on the other hand, was not influenced by material composition (**Fig.5B**). However, the addition of chitosan to the matrix showed higher ALP secretion (**Fig.5C**) and calcium deposition (**Fig.5D**). Together these results demonstrate the osteogenicity of the microgels and their material constituents.

**Figure 5:**
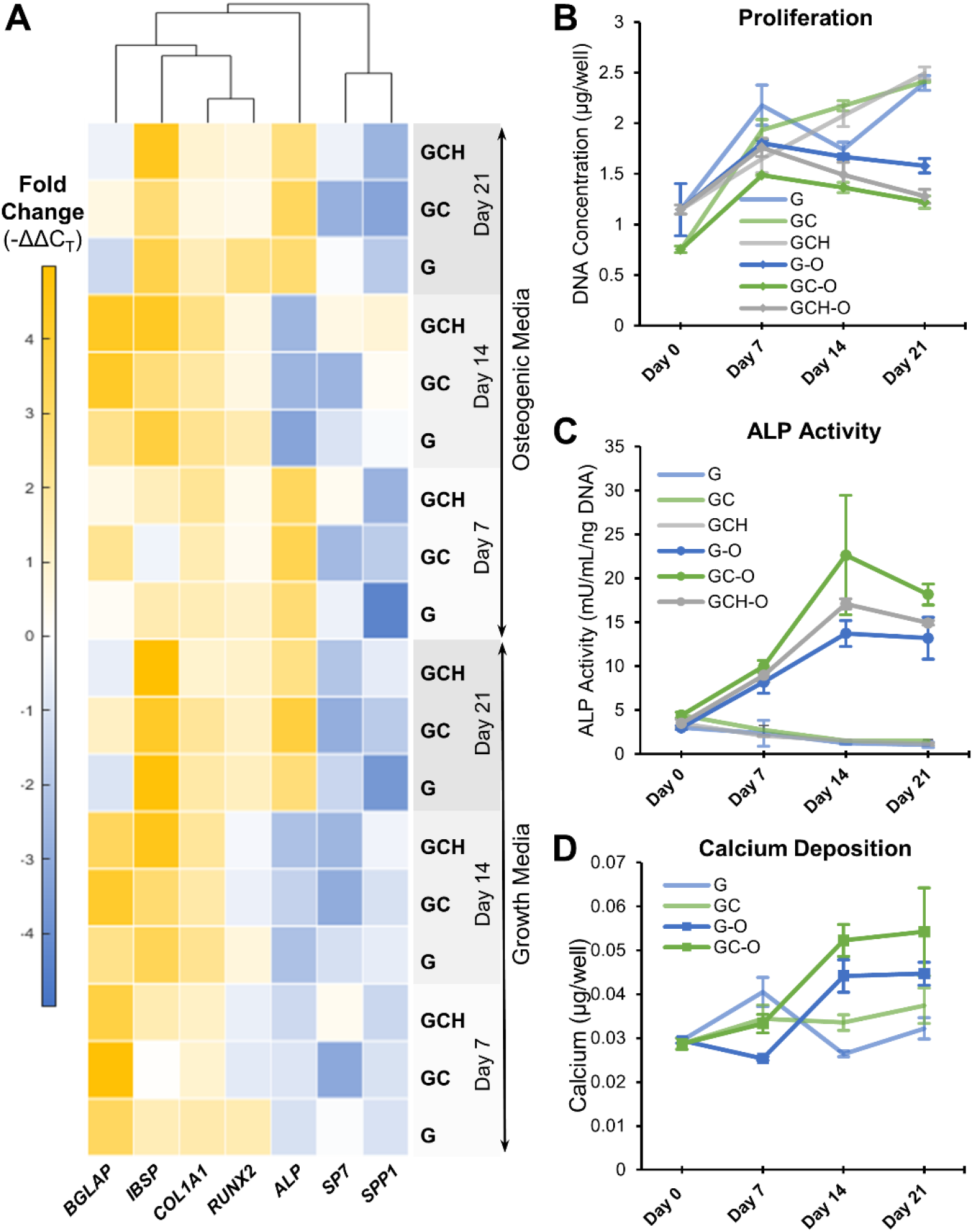
Osteogenicity of microgels with chitosan and hydroxyapatite. **A)** Osteogenic gene expression of MSC seeded in microgels. The osteogenic genes analyzed include osteocalcin (BGALP), bone sialoprotein (IBSP), collagen type I (COL1A1), Runt-related transcription factor 2 (RUNX2), alkaline phosphatase (ALP), osterix (SP7), and osteopontin (SPP1). **B)** MSC proliferation deduced from total DNA content, **C**) ALP activity quantified using a commercial kit, **D)** calcium deposition quantified using OCPC assay. GCH-group excluded due to very high background from the added HA (n = 3). Bar chart legend: G-Gelatin only, GC-Gelatin+0.25% Chitosan, GCH-Gelatin +0.25% Chitosan+1%HA. The suffix -O indicates osteogenic media.

### 3.4. Gelatin microgels support endothelial sprouting and network formation

To vascularize the constructs, we investigated the vasculogenic potential of unmodified gelatin microgels by seeding them with EC and FB. Both the cell types attached and spread well on the gelatin surface within an hour of seeding (**Fig. 6A, 6B**). The EC formed a typical ‘cobblestone’ monolayer pattern which was interrupted when seeded along with FB. The corresponding electron micrographs of the microgels show the cells firmly attached and spread on the microgel surface (**Fig. 6B**). When embedded in a 3D fibrin matrix, the endothelial cells sprouted to form vascular networks (**Fig. 6C)** within 3-5 days. The vessel density per microgel (**Fig. 6D**), the average length of a vessel segment (**Fig. 6E**), and the total vessel length (**Fig. 6F**) increased rapidly by day 10 but plateaued around day 15.

**Figure 6:**
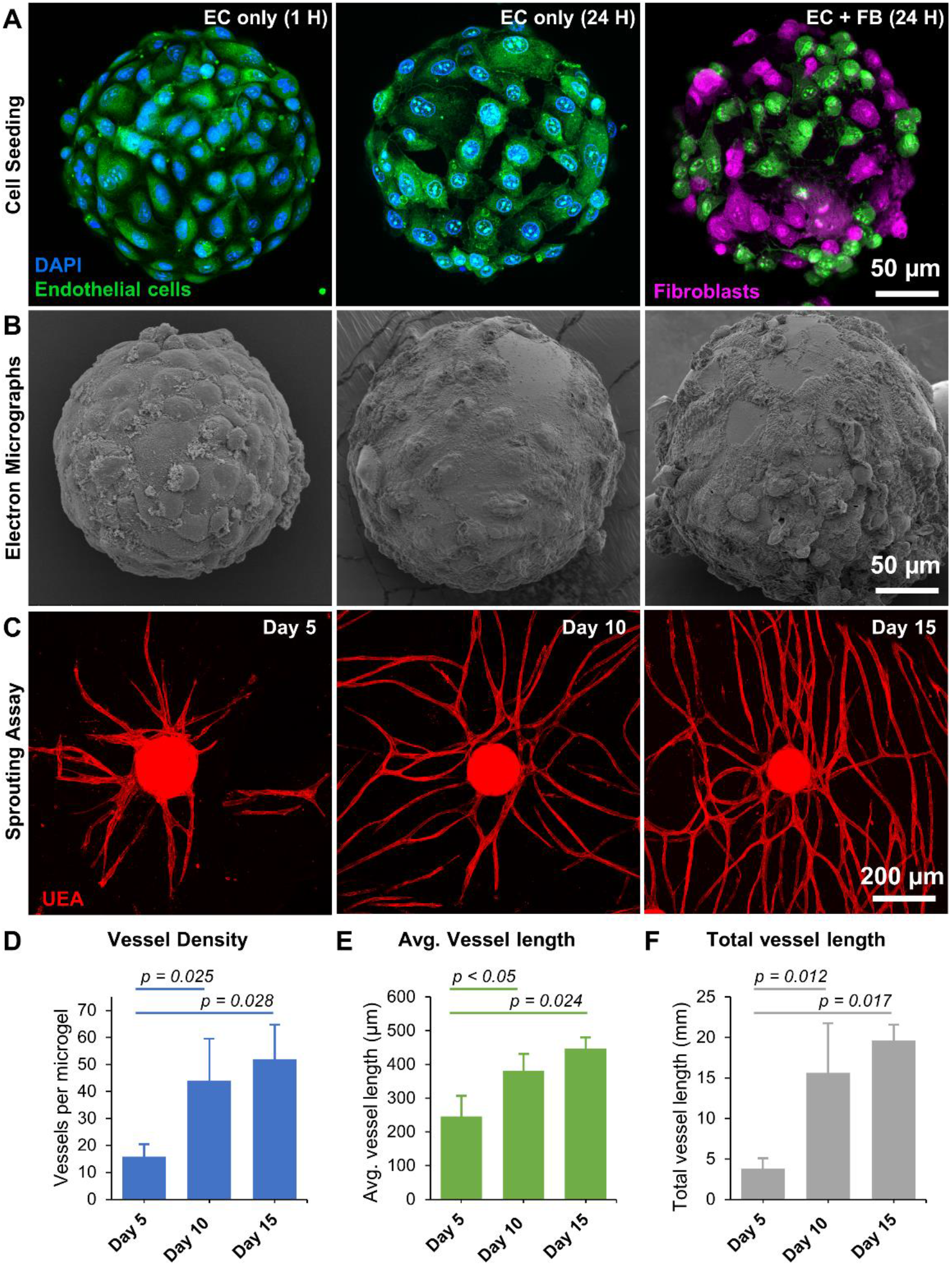
Gelatin microgels supported robust endothelial sprouting. **A)** Confocal stacks of endothelial cells and fibroblasts seeded on gelatin microgels. The endothelial cells exhibit a typical ‘cobblestone’ monolayer pattern (left and center), which was changed when seeded with fibroblasts (right). **B)** Corresponding scanning electron micrographs of microgels showing cell attachment to the microgel surface. **C)** Endothelial sprouts radiating from the microgels formed vascular networks when embedded in a fibrin matrix. **D)** Vessel density of microgel over time. **E)** Average length of a vessel segment over time. **F)** Total vessel length over time (n = 3).

The endothelial sprouts from each microgel grew radially and anastomosed with that of the neighboring microgels to form a diffuse vascular network (**Fig.7A, 7B**). The vascular anastomoses between the neighboring microgels can be seen at higher magnification (**Fig.7C, 7D**). Overall, the gelatin microgels provide a suitable microenvironment for EC and FB attachment and enable endothelial sprouting and network formation when embedded in a suitable 3D matrix.

**Figure 7.**
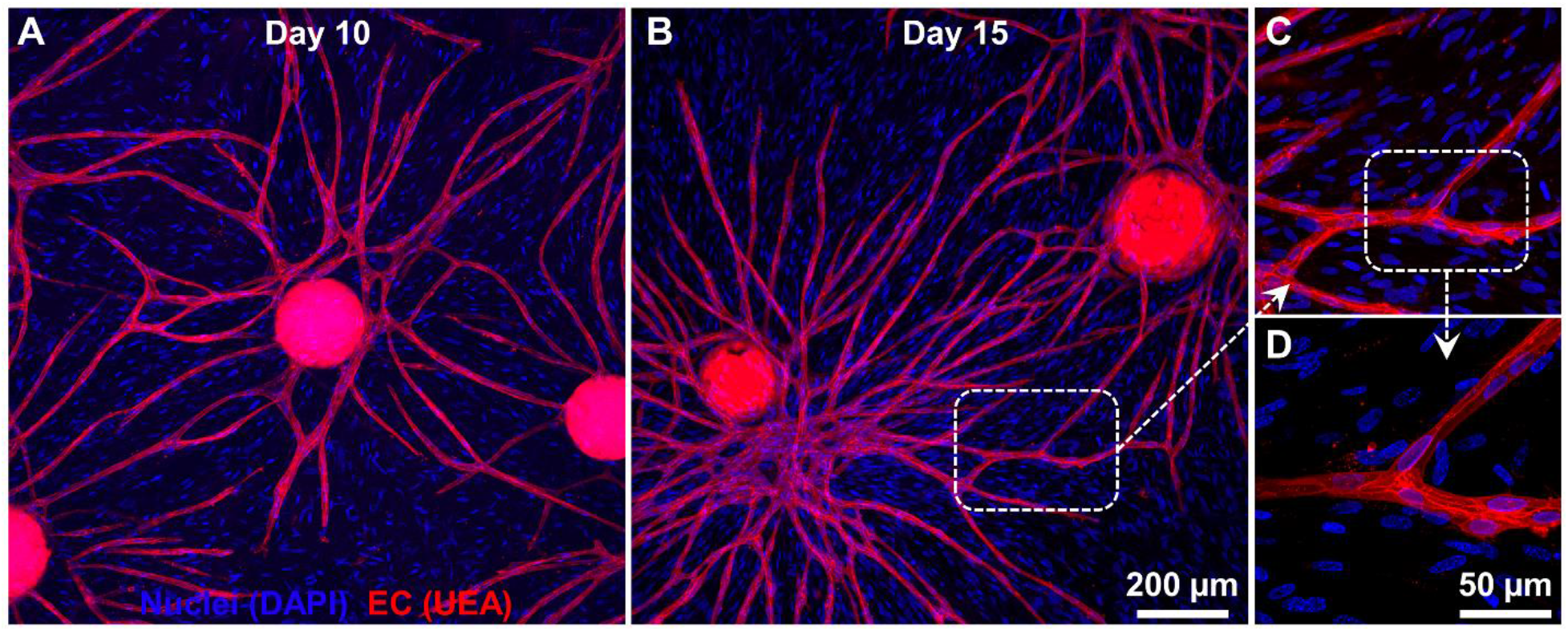
Vascular anastomoses and endothelial network formation. **A)** Confocal stacks showing the endothelial sprouts from microgel at day 10. The endothelial cells are stained with Ulex Europaeus Agglutinin I (UEA), and nuclei are stained with DAPI. **B)** By Day 15, the endothelial sprouts grew radially and anastomosed with the neighboring microgels to form a diffuse vascular network. **C** and **D)** Higher magnification confocal stacks showing vascular anastomoses between the neighboring microgels. The nuclei of the perivascular fibroblasts were visible with DAPI blue stain.

### 3.5. Creating vascularized constructs by combining osteogenic and vasculogenic microgels

To create a vascularized construct, MSC-laden osteogenic (OST) and EC/FB-laden vasculogenic (VAS) microgels were combined to form hybrid constructs and embedded in a 3D fibrin gel for characterization. The hybrid constructs were cultured in growth (GM) and osteogenic media (ODM) for 14 days. As expected, within a few days, the endothelial cells in the vasculogenic microgels sprouted and, by day 14, formed interconnected vascular networks (**Fig.8A**). The MSC seeded in the osteogenic microgels were also seen migrating in the 3D fibrin matrix by day 14 (**Fig.8A**). The total DNA content of the samples showed that the cells were actively proliferating, which plateaued after week 2 (**Fig.8B**). On the other hand, the ALP activity was higher in hybrid constructs than that of the control microgels in both growth and osteogenic media, indicating the osteogenic differentiation of MSC (**Fig.8C**). However, the total collagen matrix deposition at day 21 in the hybrid constructs was higher when cultured with osteogenic supplements (**Fig.8D**). Overall, the hybrid microgels sustained the functions of both osteogenic and vasculogenic microgels and even enhanced one another in some instances.

**Figure 8.**
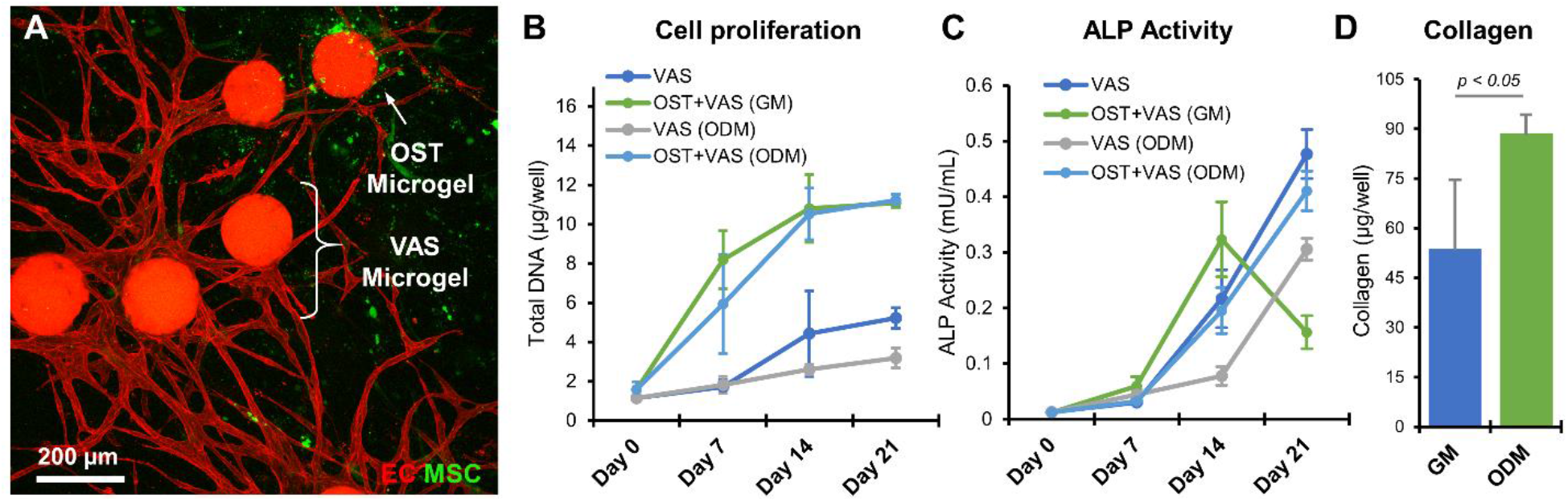
Hybrid constructs containing osteogenic and vasculogenic microgels. **A)** Confocal stacks showing the endothelial sprouts (red - Ulex Europaeus Agglutinin I) and MSC (green-CellTracker^™^ Green) in hybrid microgel constructs on day 14. **B)** Cell proliferation quantified using the total DNA content of samples at various time points. **C)** ALP activity over time. **D)** Total collagen deposition at day 21. (n = 3) Bar chart legend: OST-osteogenic microgel, VAS-vasculogenic microgels, GM-growth media, and ODM-osteogenic media.

### 3.6. Injectable microgels preserve cell viability

To demonstrate the minimally invasive application of the microgels, they were injected through a 16-25 gauge hypodermic needle, and the injection force was measured using an Instron machine equipped with a 10N load cell (**Fig. 9A**). The injection force measurements of acellular microgels showed no significant change compared to that of the PBS controls (**Fig. 9B**). Then to characterize the viability of the cell-laden microgels subjected to injection force, they were concentrated in a 1 mL syringe and injected through a 16 gauge needle and collected in a vial, and characterized for cell viability (**Fig. 9A, Inset**). The viability of the cells 1 hour and 24 hours post-injection showed no significant change in viability compared to non-injected controls (**Fig. 9C**). This shows the microgels are injectable and can be used to conformally fill complex defects.

**Figure 9.**
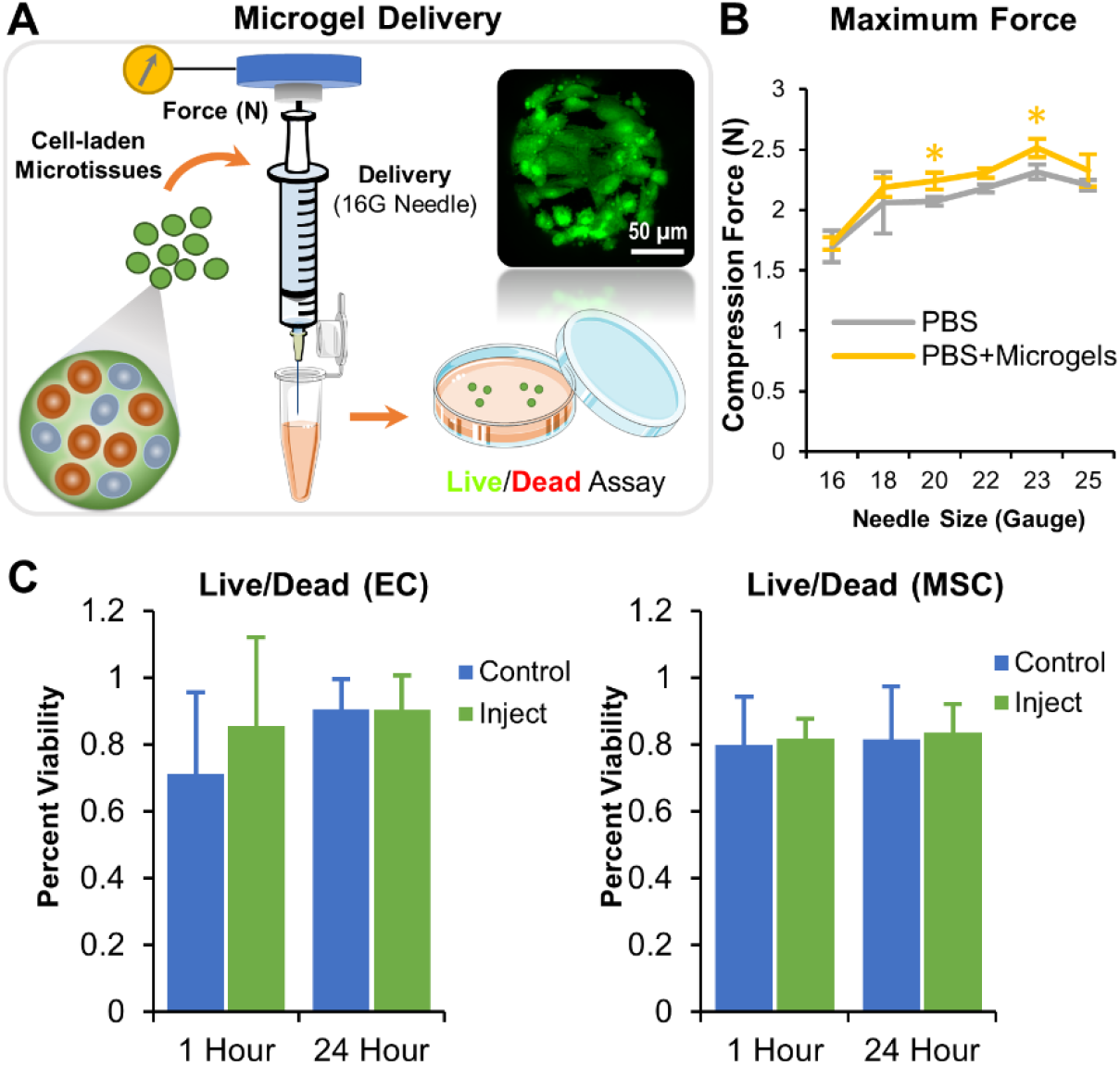
Injectability of cell-laden microgels. **A)** Test of injectability for microgels. The injection force measurement setup containing an Instron Universal Testing machine equipped with a 10 N load cell measured the compression force. (Inset micrograph) Live human umbilical vein endothelial cells, stained with calcein AM, on the surface of a microgel that was passed through a 16 gauge needle (n = 5). **B)** The injection force measurements of acellular microgels using a hypodermic needle of various sizes showed no significant difference compared to the PBS controls. *p = 0.013 (20 gauge) and 0.003 (23 gauge). **C)** The viability of the EC and MSC, 1 hour and 24 hours post-injection, showed no significant change in viability compared to non-injected controls (n = 12).

### 3.7. Bone formation by osteogenic microgels in critical-sized defect

The ability of microgels to regenerate bone was then evaluated in an established murine full-thickness critical-sized cranial defect model. A 3 mm wide nonsuture-associated parietal bone defect on an adult mouse (>12 weeks old) is shown to not heal spontaneously during the lifetime of the animal^35^. Pure gelatin microgels and gelatin microgels with 0.25% chitosan, 1% HA were fabricated, seeded with murine MSC (1.0×10^6^ cells/200 μL of microgels), and cultured under osteogenic conditions for 12 days. A fibrin carrier gel was then used to concentrate the microgels and contain them within the critical-sized defect in mice. Animals that received acellular fibrin carrier gel aline served as control. Gross examination of the 3D reconstructed microCT images showed new bone formation within the defect resulting in 0%, ~65% and >95% defect closure by week 12 in control, microgel, and microgel with HA groups, respectively (**Fig. 10A**). Bone morphometric analysis showed a steady increase in new bone volume and bone mineral density (BMD) of the construct over 12 weeks in both microgel conditions (**Fig. 10B**). At week 12, a total of 0.25±0.18 mm^3^ of new bone volume and 0.24±0.12 mg of HA/mm^3^ mineral density was seen in microgels (**Fig. 10B**). A significantly higher total new bone volume of 0.7±0.28 mm^3^ and a mineral density of 0.34±0.09 mg of HA/mm^3^ were seen in HA microgels. No noticeable new bone formation was seen in control conditions. A detailed statistical comparison of bone morphometry parameters between conditions is listed in **Table.2**.

**Figure 10.**
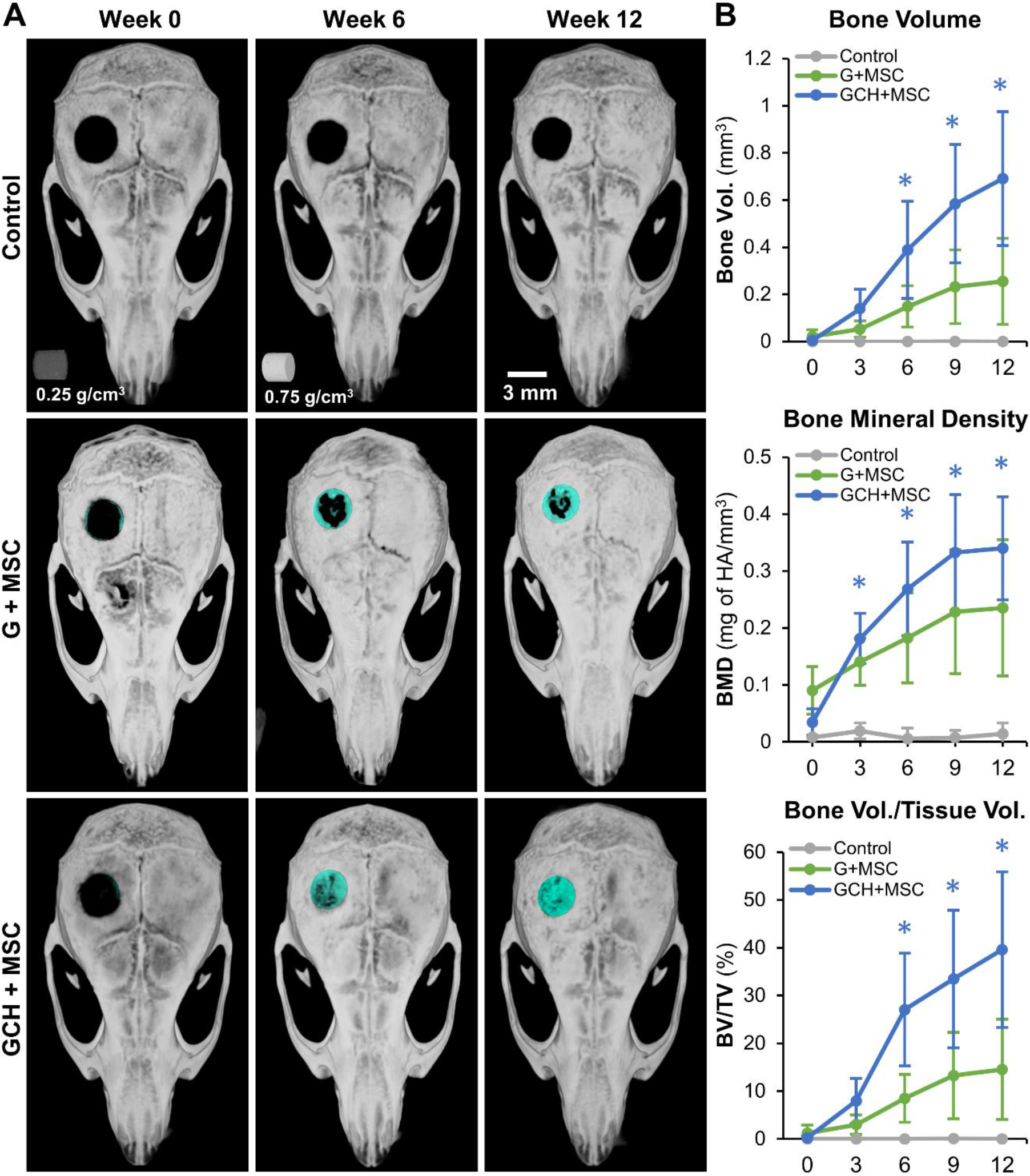
Murine critical-sized calvarial defect. **A)** 3D reconstructed microCT images of the mice crania over time, showing the defect being bridged. Control (CNT) animals received acellular fibrin carrier gel, the second group received gelatin microgels seeded with MSC, and the last group received gelatin microgels with HA seeded with MSC. **B**) Bone morphometric analysis of the newly formed bone shows a steady increase in new bone volume and mineral density (BMD) in conditions containing MSC-seeded microgels. n = 4 for each condition.

**Table 2:** Holm-Sidak comparison of bone morphometry parameters between conditions. (*Control* = *No microgel control, G*+*MSC* = *Gelatin microgels seeded with MSC, GCH*+*MSC* = *Gelatin microgels with chitosan and HA seeded with MSC*.)

| Time | Comparison | Bone Vol.<br>(*p*-value) | BMD<br>(*p*-value) | BV/TV<br>(*p*-value) |
| --- | --- | --- | --- | --- |
| **Week 0** | G+MSC vs Control | 0.995 | 0.335 | 0.995 |
| **Week 0** | GCH+MSC vs Control | 0.971 | 0.64 | 0.971 |
| **Week 0** | GCH+MSC vs G+MSC | 0.981 | 0.506 | 0.981 |
| **Week 3** | G+MSC vs Control | 0.601 | 0.053 | 0.601 |
| **Week 3** | GCH+MSC vs Control | 0.485 | 0.02 | 0.484 |
| **Week 3** | GCH+MSC vs G+MSC | 0.626 | 0.441 | 0.625 |
| **Week 6** | G+MSC vs Control | 0.144 | 0.004 | 0.144 |
| **Week 6** | GCH+MSC vs Control | 0.002 | <0.001 | 0.002 |
| **Week 6** | GCH+MSC vs G+MSC | 0.041 | 0.111 | 0.04 |
| **Week 9** | G+MSC vs Control | 0.026 | <0.001 | 0.026 |
| **Week 9** | GCH+MSC vs Control | <0.001 | <0.001 | <0.001 |
| **Week 9** | GCH+MSC vs G+MSC | 0.002 | 0.055 | 0.002 |
| **Week 12** | G+MSC vs Control | 0.015 | <0.001 | 0.015 |
| **Week 12** | GCH+MSC vs Control | <0.001 | <0.001 | <0.001 |
| **Week 12** | GCH+MSC vs G+MSC | <0.001 | 0.054 | <0.001 |

Histological analysis of the demineralized skulls shows control condition treatment with no bridging, while the gelatin microgels seeded with MSC showing partial bridging with collagen and elastic fibers (**Fig. 11**). The gelatin-chitosan-HA microgels exhibited the highest bridging with the osteoid formation and collagenous matrix deposition (**Fig. 11**). Together, the osteogenic microgels enabled MSC to bridge the critical-sized defect with mineralized collagen-rich osteoid bone.

**Figure 11.**
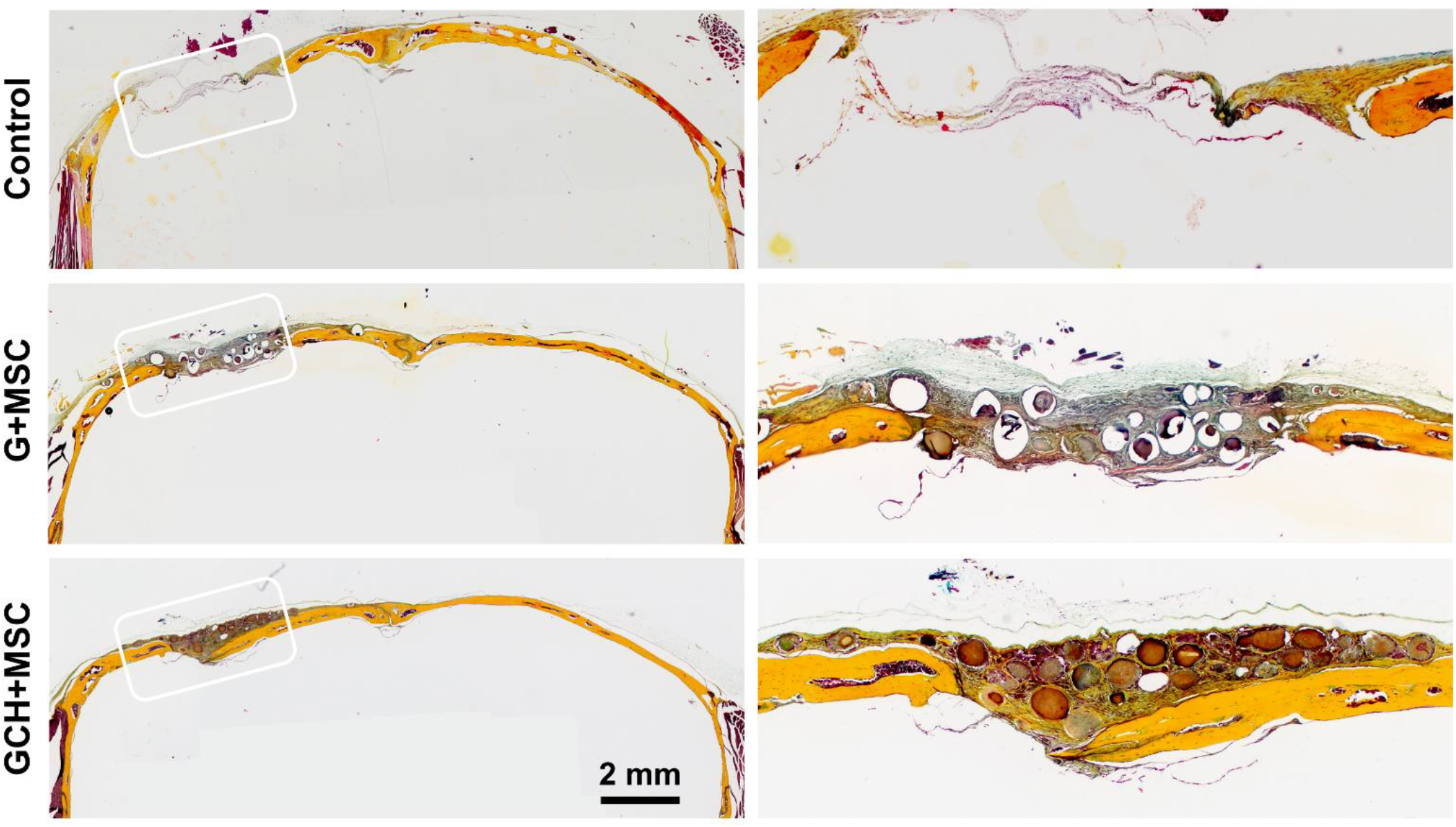
Gelatin microgels with MSC regenerate bone in a critical-sized calvarial defect. Movat’s Pentachrome staining of the demineralized bone sections shows healing of the critical-sized defect in various treatment conditions. A yellow stain indicates collagen fibers, and a black stain indicates elastic fibers. (Left) The control condition showed no bridging, while the gelatin microgels seeded with MSC showed partial bridging with collagen and elastic fibers. The gelatin-chitosan-HA microgels exhibited the highest bridging with the osteoid formation and collagenous matrix deposition. (Right) Magnified images of the defect area of corresponding treatments shown on the left.

## 4. Discussion

The hypo-immunogenicity of MSC and their ability to self-renew and differentiate into bone, cartilage, and adipose tissues have made them an attractive adult source for tissue engineering. The MSC are also known for their immunomodulatory and angiogenic factor secretion, further enriching the regenerative environment^36^. Here we investigated the application of an off-the-shelf nanoporous microgel for MSC-based bone tissue engineering.

### 4.1. Tunable microgels for natural materials

The microgel’s primary constituent is gelatin, a well-characterized material^37, 38^ consisting of water-soluble polypeptides derived from partially denatured collagen. The polypeptides have integrin-binding sites as parent collagen^39^ but are less immunogenic due to the removal of aromatic groups^40^. Gelatin has a sol-gel transition near 30°C. Hence, to stabilize them in physiological conditions, it is necessary to crosslink the matrix, which also helps integrate other materials such as chitosan into the matrix. The method and extent of the crosslinking immensely influence the mechanical properties, swelling ratio, cytotoxicity, antigenicity, and rate of degradation of the hydrogel^26, 37^. Chemical crosslinkers such as formaldehyde^41^, glutaraldehyde^42^, and glyceraldehyde^43^ are commonly used to stabilize gelatin matrix and enhance resistance to thermal degradation, but their cytotoxicity is a significant concern. We used a naturally occurring crosslinking agent, genipin, obtained from the fruits of *Gardenia jasminoides*. They are significantly less cytotoxic and genotoxic^44^ with an intravenous LD_50_ of 153 mg/kg^45^, which is 10 fold higher than glutaraldehyde (mouse LD_50_15.4 mg/kg i.v.). The genipin-crosslinked gelatin is enzymatically degradable, and the rate is concentration dependant^29^. Further, genipin crosslinking allows controlling the degree of crosslinking through concentration and incubation time^46^.

We tested the propeller and parallel-plate impeller (aka flat-blade turbine) for emulsification to fabricate the microgels. Both are well-suited for the 100 cS (0.1 kg/m-s) PDMS used, but they create axial and radial flow patterns, respectively^47^. The parallel-plate impeller had a bimodal size distribution due to solid body rotation and limited liquid mixing above and below the impeller. The parallel-plate impeller is efficient in power consumption and dispersion, but enhancements such as baffles may be necessary to avoid solid body rotation and a large central surface vortex. On the other hand, the axial flow pattern created by the propeller impeller yielded microgels with sizes showing normal distribution and a linear decrease in mean and median values with an increase in mixing speed. Further, the desired size range of the microgels was achieved with less turbulence in the mixing vessel. Hence the propeller impeller yielded the best outcomes for the microgel fabrication and was therefore chosen for the rest of the studies.

### 4.2. Microgels create a conducive environment for osteogenesis and vascularization

The material constituents and physical parameters of the microgel greatly influence the osteogenicity of the microenvironment. Tissue softness is a “colligative” property as the water content of the tissue is inversely proportional to the tissue elasticity and collagen content^48^. The average bone matrix hydration is around 40% compared to that of the brain, which is around 90%. The nanoporous structure and the high polymer density of the microgels arising from the highly cross-linked polypeptides (~90% degree of crosslinking) enable the matrix to hold large amounts of bound water and swell ~5 fold in volume. The modest water content and high collagen density give rise to the stiffness (0.1 - 30 MPa) and rigidity of the bone, which subsequently regulate bone function. Even during fractures, mechanobiological studies point to the mechanics of the callus (specific stiffness range of 0.1-1 MPa) to be a crucial promotor of osteogenic differentiation of progenitor cells and neovascularization^49^. Similarly, a stiffness of ~100 kPa is shown to regulate endothelial progenitor cell differentiation and vascularization^49, 50^. Hence, the measured stiffness of hydrated microgels (<200 kPa) falls in a range that is conducive to both osteogenesis and vasculogenesis.

Further, in the body, MSC undergo osteogenesis when attached on the surface of the stiff bone with the soft marrow above. The 2D surfaces appropriately mimic the process of MSC on the bone surface and differentiation into osteoblasts^51^. The microgels with a high polymer density, moderate stiffness, and modest hydration show a similar ability to support MSC attachment, proliferation, and osteogenesis. To enhance their osteogenicity, we tailored the formulation to mirror the native bone matrix, which increases osteogenic differentiation in MSC^52, 53^. When we supplemented the microgel matrix with chitosan and HA, it enhanced MSC differentiation even while retaining the properties of the pure gelatin microgels. Apart from material composition, the osteogenicity of the microgels can also be attributed to their geometry. A micron-scale curvature is shown to naturally induce cytoskeletal stress, which leads to the osteogenic differentiation of MSC via RhoA/ROCK pathways^19, 54, 55^.

### 4.3. Dual-phase constructs promote synergism

Aside from the parenchymal cells, vessel growth and blood flow are vital in determining bone repair and osteogenesis. Distinct capillary subtypes in the bone are known to secrete angiocrine and osteogenic factors and maintain osteoprogenitors^56^. Vessel growth and osteogenesis are coupled through molecular crosstalk between endothelial cells and osteoblasts^57^. To harness this synergistic association, we mixed the osteogenic microgels with vasculogenic microgels containing endothelial cells and pericytes. Typically, cultured endothelial cells quickly lose their ability to adopt tissue-specific features required for organ development and regeneration. Although, this adaptability may be revived with a suitable blend of matrix proteins^58^. We show that endothelial cells seeded in gelatin-only microgels retain their ability to sprout and form vascular plexi. But they might need an appropriate pericyte-like cell for providing the growth factor as robust sprouting was absent when endothelial cells were seeded alone. Similarly, the endothelial cells failed to sprout when the fibroblasts were seeded in the embedding matrix instead of the microgel (data not shown). Seemingly the pericytes also function better when provided with the support of the microgel, although more mechanistic studies are warranted to elucidate these effects. When we combined the vasculogenic microgels with osteogenic ones, they sustained endothelial sprouting while supporting the osteogenic phenotype. Further, the dual-phase constructs provide the flexibility to optimize formulations and culture mature the components independently without limiting each other. At the same time, they can be easily combined when necessary to form a single homogenous construct that can promote vascularized tissue regeneration. These dual-phasic constructs are valuable, especially to heal complex fractures that are often ischemic and require concurrent vascularization.

### 4.4. Injectable microgels promote bone regeneration

Minimally invasive applications can reduce surgery-related complications, including infection and long recovery times. Further, the microscale modules allow them to be injectable and conformally fill complex defects^14^. When we injected our microgels using a hypodermic needle of varying sizes, we saw no change in compression force or clogging of microgels. The shear-induced damage to the cells attached to the surface is also negligible. These injectable microgels are an off-the-shelf alternative to the existing bone grafts. When combined with autologous blood, plasma, or bone marrow, their very high swelling ratios can enable them to quickly soak in growth factors while attracting adherent mesenchymal progenitor cells onto their surface. In addition, carrier gels such as fibrin glue can also be used to concentrate the microgels in the wound site^14^.

To validate the ability of microgels in regenerating bone, we used a murine calvarial defect model. Although our calvarial model generates a critical-sized defect, it doesn’t create an ischemic wound. So, we validated the osteogenic gelatin microgels alone without a vasculogenic component. As defined by the FDA, a nonunion is a fracture that persists for a minimum of nine months without any signs of healing for three months^59^. We show that our implanted microgels supported robust bone regeneration and showed continual progress with 65-95% defect closure by week 12, depending on the microgel formulation. As hypothesized, the addition of chitosan and HA to the gelatin matrix significantly accelerated the new bone formation and defect closure. In addition, histological analysis of explants confirmed the effectiveness of the osteogenic microgel in regenerating bone with a rich collagenous matrix indicative of robust osteogenic differentiation. Overall, our data demonstrate the therapeutic ability of the osteogenic microgels, and further studies on an ischemic defect model are warranted to validate concurrent vascularization.

## 5. Conclusion

Together, our work shows that microgels can be used to augment the differentiated function of MSC and endothelial cells to fabricate a vascularized bone construct. The microgel matrix is customizable to mimic tissue-specific functions while allowing minimally invasive delivery. Microgels support robust bone regeneration and healing of critical-sized defects, as evident from our mouse studies. Our work lays the foundation to establish principles of designing multiphasic scaffolds with tissue-specific biophysical and biochemical properties for regenerating vascularized and interfacial tissues.

## 6. Acknowledgments

Research reported in this publication was supported in part by the National Institute of Arthritis and Musculoskeletal and Skin Diseases (NIAMS) Award Number R21AR078447, National Institute of General Medical Sciences of the National Institutes of Health under Award Numbers P20GM130456 and P20GM103436-20 (KY IDeA Networks of Biomedical Research Excellence), National Center for Research Resources and the National Center for Advancing Translational Sciences of the National Institutes of Health under Award Number UL1TR001998, and Orthopedic Trauma Association (OTA, Grant Number: 6889). The content is solely the responsibility of the authors and does not necessarily represent the official views of the National Institutes of Health or other grant funding agencies.

## 7. Data Availability

All data generated or analyzed during this study are included in this published article.

